# Gain of 1q confers an MDM4-driven growth advantage to undifferentiated and differentiating hESC while altering their differentiation capacity

**DOI:** 10.1101/2023.09.19.558389

**Authors:** Nuša Krivec, Edouard Couvreu de Deckersberg, Yingnan Lei, Diana Al Delbany, Marius Regin, Stefaan Verhulst, Leo A. van Grunsven, Karen Sermon, Claudia Spits

## Abstract

Gains of 1q are a highly recurrent chromosomal abnormality in human pluripotent stem cells. In this work, we show that gains of 1q impact the differentiation capacity to derivates of the three germ layers, leading to miss-specification to cranial placode and non-neural ectoderm during neuroectoderm differentiation and by poorer expression of lineage specific markers in hepatoblasts and cardiac progenitors. Competition assays show that the cells retain their selective advantage during differentiation, which is mediated by a higher expression of *MDM4*, a gene located in the common region of gain. *MDM4* drives the winner phenotype of the mutant cells in both the undifferentiated and differentiating state by reducing the cells’ sensitivity to DNA-damage through decreased p53-mediated apoptosis. Finally, we find that cell density in culture plays a key role in promoting the competitive advantage of the cells by increasing DNA damage.

## INTRODUCTION

At present, over 7400 hPSC lines have been registered at the hPSC lines registry (https://hpscreg.eu) and over 50 ongoing and completed clinical trials involve the transplantation of cells derived from hPSC^1,2^. While the ability to differentiate is a fundamental characteristic of all hPSCs, they may differ in their differentiation capacity to specific lineages or cell types^3^. Causes for this variability are common genetic variation^4,5^, epigenetic variance^6–8^, mitochondrial mutations and a broad range of environmental factors induced by differences in culture conditions^3^. Another significant source of variation are genetic changes acquired by the cells during in vitro culture, such as copy number variations^9,10^ and single nucleotide changes such as p53-inactivating mutations^11–13^.

Full or segmental gains of chromosomes 1, 12, 17 and 20 are known to be highly recurrent in both human induced pluripotent stem cells (hiPSC) and human embryonic stem cells (hESC)^9,10,14^. They confer the cells increased cloning efficiency, decreased doubling times, decreased sensitivity to apoptotic triggers, increased self-renewal and growth factor independence^15–19^. This results in a growth advantage in the undifferentiated state and a rapid culture take-over by the mutant cells^17,20^. Presently, the mechanisms and driver gene behind this selective advantage has been fully characterized for only one of these recurrent abnormalities: the gain of 20q11.21 results in a decreased sensitivity to apoptosis due to higher *BCL2L1* expression^16,17^. Whether this selective advantage is maintained during differentiation remains unanswered. Further, there is increasing evidence that these recurrent abnormalities affect the differentiation capacity of hPSC, often in a cell-lineage specific manner^15,18,21^. Gains of 20q11.21 and losses of 18q lead to impaired neuroectoderm commitment^22,23^ and gain in chromosome 12p results in large foci of residual undifferentiated cells persisting after hepatoblast differentiation^24^.

In this study, we focused on the gain of 1q, which is one of the most commonly acquired genetic abnormalities in hPSC worldwide^9^, while little is known on its functional effects. Human ESC with a complex karyotype, including a gain of 1p, have an altered gene expression profile with an activation of WNT signaling and deregulation of tumor suppressors and oncogenes, as well as a differentiation bias towards ectoderm^21^ and gain of 1q was found to recurrently appear during in vitro neural differentiation, suggesting a selective advantage during differentiation^25^. In this work, we found that gain of 1q alters the differentiation capacity of hESC and established that *MDM4*, a regulator of p53 located in the minimal region of gain, drives their competitive advantage both in the undifferentiated state and during differentiation by reducing their sensitivity to DNA-damage induced p53-mediated apoptosis.

## RESULTS

Figure 1 shows the overall setup of this study. All experiments were carried out on the hESC lines VUB03 and VUB19^26,27^. In vitro culture led to three independent events of gain in chromosome 1. VUB19 and one subline of VUB03 acquired a gain of the entire q arm of chromosome 1 (termed VUB19^1q21.1qter^ and VUB03^1q21.1qter^), another subline of VUB03 gained a smaller region spanning 3.3 Mb in 1q32.1 (VUB03^1q32.1^).

### hESC^1q^ mis-specify to placode and non-neural ectoderm during neuroectoderm differentiation and differentiate to immature hepatoblasts and cardiac progenitors than hESC^wt^

We investigated the impact of the gain of 1q on trilineage differentiation by subjecting hESC^wt^ and hESC^1q^ to neuroectoderm (NE), hepatoblast (HEP) and cardiac progenitor (CP) differentiation (VUB03^1q21.1qter^, VUB19^1q21.1qter^, VUB03^1q32.1^, VUB03^wt^ and VUB19^wt^, all lines differentiated at least in triplicate, Figure 1). We measured the expression of six lineage-specific markers and of *NANOG* and *POUF51* to evaluate differentiation efficiency and to test for residual undifferentiated cells (Figure 2A and 2C and Figure S1). All differentiated cells had almost undetectable levels of *POU5F1* and *NANOG* mRNA, and no POU5F1-positive cells appeared in the immunostainings (Figure 2A), indicating that all cells exited the undifferentiated state (Figure 2C and Figure S1).

**Figure 1:**
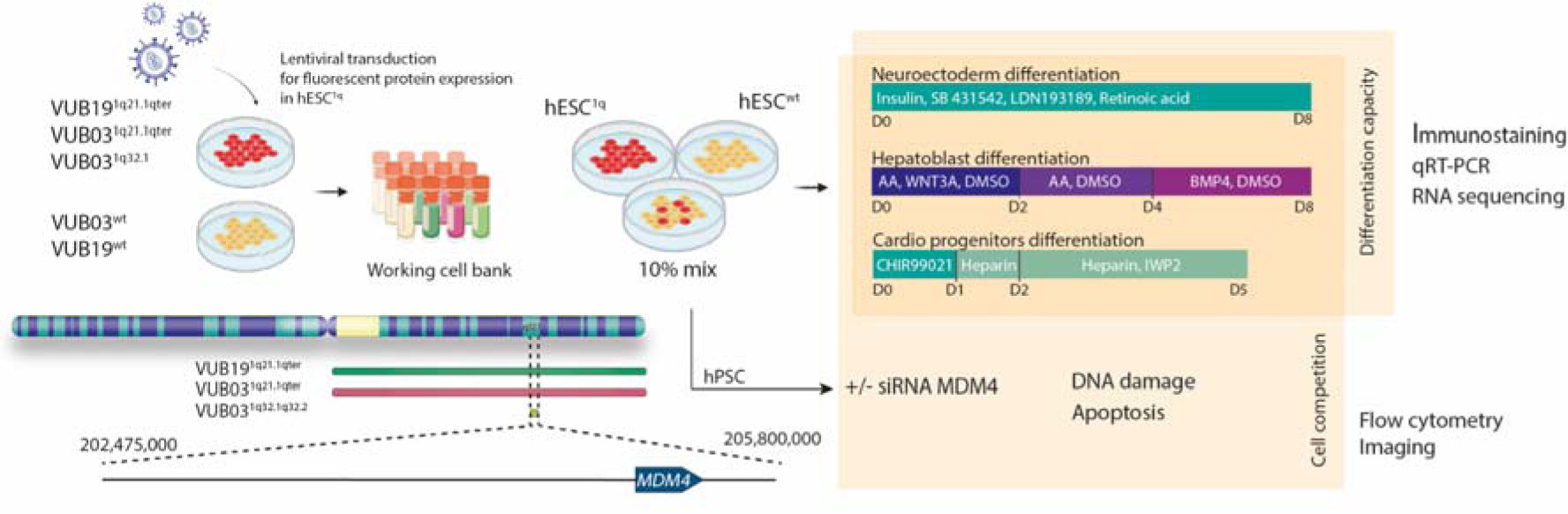
Schematic representation of experimental setup of this study. Lentiviral transduction was used to induce stable expression of fluorescent proteins in hESC^1q^. Working cell banks were established comprising mutant labeled cell lines and their isogenic counterparts. Pure wild-type and 1q lines were utilized, along with mixes containing 10% of 1q cells, for differentiation to three lineages: neuroectoderm, hepatoblasts and cardiac progenitors. Differentiation capacity was assessed through immunostaining, qPCR, and RNA sequencing. Cell competition results were measured by analyzing the ratio of mutant and wild-type cells before and after differentiation using flow cytometry and immunostaining. To study the role of *MDM4* in the growth advantage of hESC^1q^, *MDM4* was downregulated by siRNA prior to the competition assays. To study the mechanisms by which *MDM4* provides the selective advantage, DNA damage was induced by bleomycin and apoptosis levels were evaluated using Annexin V staining and flow cytometry, while DNA damage levels were studied through immunostaining for gamma-H2AX.

**Figure 2.**
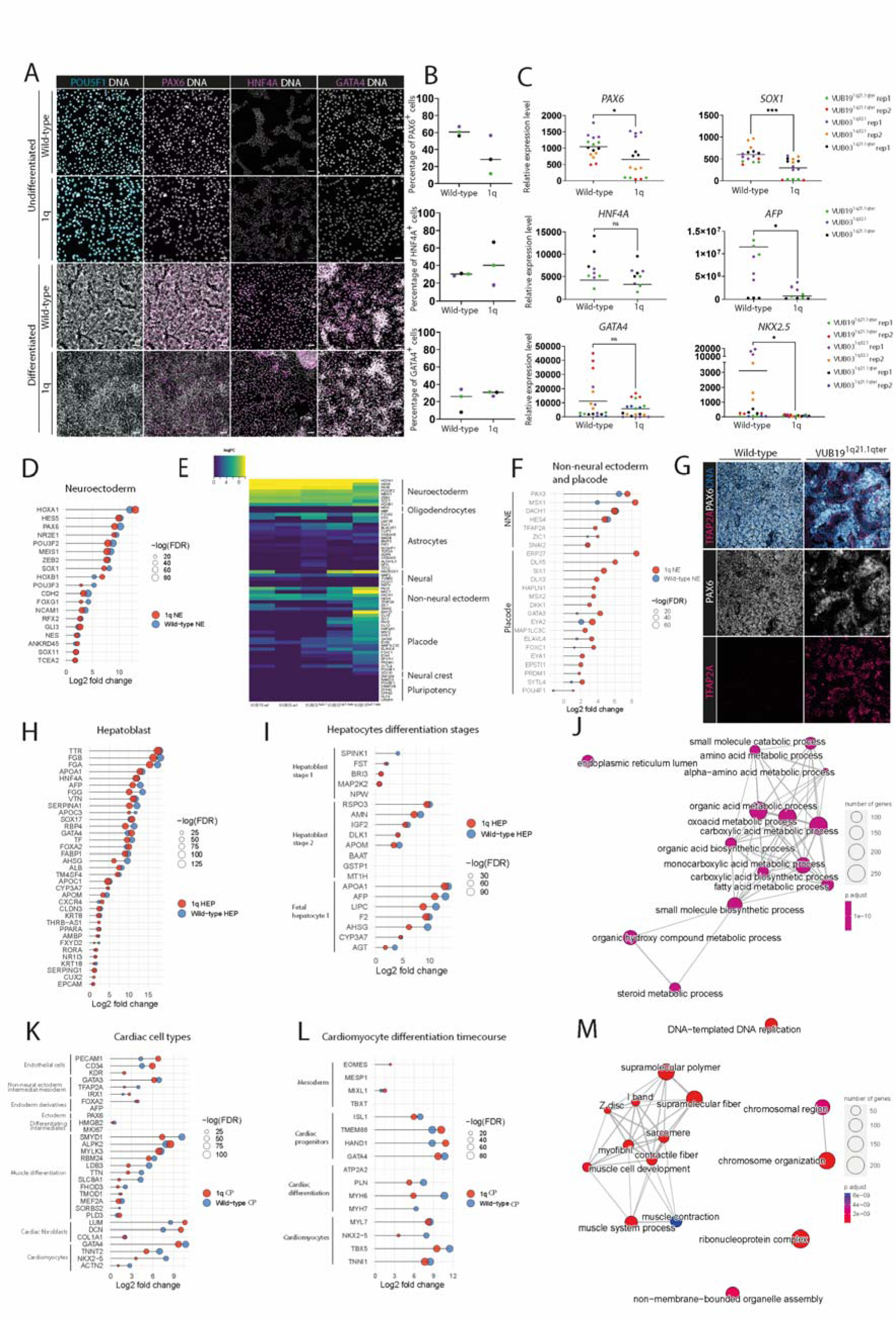
**hESC^1q^ miss-specify to placode and non-neural ectoderm during neuroectoderm differentiation and differentiate to more immature hepatoblasts and cardiac progenitors than hESC^wt^.** A Examples of immunostaining for POU5F1, PAX6, HNF4A and GATA4 of hESC^wt^ and hESC^1q^ before and after the completion of induction of differentiation to neuroectoderm (NE, day 8), hepatoblast (HEP, day 8), cardiac progenitors (CP, day 5) of VUB19^1q21.1qter^. Figure 1S shows the images for cell lines VUB03^1q21.1qter^ and VUB03^1q32.1^ included in the study. B Quantification of the number of PAX6, HNF4A and GATA4-positive cells in the immunostainings shown in A and in Figure 1S. No statistically significant differences were found. C mRNA quantification by quantitative real-time PCR of *PAX6* and *SOX1* in NE^wt^ and NE^1q^, *HNF4A* and *GATA4* in HEP^wt^ and HEP^1q^, and *GATA4* and *NKX2.5* in CP^wt^ and CP^1q^; * p<0.05, *** p<0.001. Results for *POU5F1* and NANOG are shown in Figure 1S. Each differentiation was carried out in three technical replicates at the same time. Different hESC lines and independent experimental replicates (rep) are coded in different colors. D Lollipop diagrams representing the differential gene expression of neuroectoderm markers in NE^wt^ and NE^1q^ vs undifferentiated cells. All genes shown in this panel and in E, H, I, K, and L are induced with a log_2_fold-change over 0 and FDR<0.05. E Lollipop diagrams representing the differential gene expression of placode markers in NE^wt^ and NE^1q^ vs undifferentiated cells. F Heatmap representing the induction of expression of genes marking different ectodermal cell types and the undifferentiated hESC state in NE obtained from the five lines included in this study. G Representative images of the immunostaining of NE^wt^ and NE^1q^ for PAX6 and TFAP2A. H Lollipop diagrams representing the differential gene expression of hepatoblast markers in HEP^wt^ and HEP^1q^ vs undifferentiated cells. I Lollipop diagrams representing the differential gene expression of markers of different stages of hepatocyte differentiation in HEP^wt^ and HEP^1q^ vs undifferentiated cells. J Enrichment map showing the top-15 deregulated pathways from gene ontology gene set enrichment analysis in HEP^wt^ vs HEP^1q^. The size of nodes indicates the number of genes in each pathway and the color represents adjusted p-value. Pathways that cluster together have overlapping gene sets. K Lollipop diagrams representing the differential gene expression of genes marking different cardiac cell types in CP^wt^ and CP^1q^ vs undifferentiated cells. L Lollipop diagrams representing the differential gene expression of genes marking different stages of cardiomycte differentiation in CP^wt^ and CP^1q^ vs undifferentiated cells. M Enrichment map showing the top-15 deregulated pathways from gene ontology gene set enrichment analysis in CP^wt^ vs CP^1q^.

NE^1q^ showed significantly lower mRNA levels of *PAX6* and *SOX1* compared to the levels in NE^wt^ (Figure 2C p*^PAX6^*=0.0323 and p*^SOX1^*=0.0002, unpaired t-test, N=15), indicating a decreased neuroectodermal differentiation efficiency. In line with this, immunostaining showed a lower percentage of PAX6-positive cells in differentiated hESC^1q^ than in their isogenic hESC^wt^ counterparts, but without reaching statistical significance (Figure 2B, NE^1q^=32.15%, NE^wt^=61.24%, p=0.0974, unpaired t-test, N=3). For the HEP differentiation, we found no significant difference between HEP^wt^ and HEP^1q^ in the mRNA expression levels of *HNF4A* while *AFP* had a significantly lower expression in HEP^1q^ (Figure 2C, p^HNF4A^=0.2247, p^AFP^=0.0142, unpaired t-test, N=9) and immunostaining for HNF4A showed similar percentages of HNF4A positive cells in both groups (Figure 2B, HEP^1q^=41.60%, HEP^wt^=29.79%, p=0.4514, unpaired t-test, N=3). Similarly, the mRNA levels of the CP marker GATA4 was not significantly different between CP^wt^ and CP^1q^ while *NKX2-5* was significantly lower in CP^1q^ (Figure 2C, p^GATA4^=0.1537, p^NKX2.5^=0.0398, unpaired t-test, N=18), which was confirmed by the immunostaining for GATA4 (Figure 2B, CP^1q^=29.56%, CP^wt^=22.74%, p=0.4390, unpaired t-test).

Taken together, these results show that while hESC^1q^ show a decreased differentiation efficiency into neuroectoderm, they do not remain undifferentiated, suggesting that part of the cells miss-specify to alternative cell fates. For the mesendoderm lineages, hESC^1q^ appear to commit equally efficiently to this germ layer than their genetically balanced isogenic counterpart, as reflected by the similar expression for the early markers of endoderm and mesoderm induction, while showing a decreased expression of the markers for further cell-type commitment.

To further characterize the differentiated cells, we carried out bulk mRNA sequencing of 43 samples: 15 samples of neuroectoderm (NE^wt^=6, NE^1q^=9, 14 of cardiac progenitors (CP^wt^=6, CP^1q^=8) and 14 of hepatoblasts (HEP^wt^=6, HEP^1q^=8). Differential gene expression analysis showed that NE^1q^ differentially expressed 1603 genes as compared to NE^wt^, while HEP^1q^ and CP^1q^ differentially expressed 189 and 241 genes respectively as compared to HEP^wt^ and CP^wt^ (Figure S2, |log_2_fold-change|>1.0 and false discovery rate (FDR)<0.05). We surmised that this significant difference in the number of differentially expressed genes was caused by miss-specification of hESC^1q^ to neuroectoderm and were rather yielding a mixed cell population, which did not occur in the mesendoderm lineages.

To determine the alternate cell fate acquired by hESC^1q^ upon neuroectoderm differentiation, we carried out differential gene expression analysis of the NE^1q^ and the NE^wt^ relative to bulk RNA sequencing data of undifferentiated hESC (N=38 samples, archived lab data). First, we tested the expression of neuroectoderm markers, which we found to be less induced in both NE^1q^ than in NE^wt^ (Figure 2D). We then queried the data for the expression of different sets of markers for embryonic and extra-embryonic lineages that appear in early human development. Table S1 shows the lists of gene sets we curated from published data^28–44^. We found NE^1q^ showed high levels of expression of markers of non-central nervous system (non-CNS) ectodermal lineages such as non-neural ectoderm and of cranial placode (Figure 2E and Figure 2F, Figure S3). To confirm this, we double-stained NE^wt^ and NE^1q^ for the non-CNS marker TFAP2A^39^ and the NE marker PAX6 and found that while hESC^wt^ efficiently differentiate to a homogeneous neuroectoderm cell-population, hESC^1q^ produce a mix of neuroectoderm and non-CNS cells (Figure 2G).

Next, we used this same approach to study the cell types obtained from the HEP and CP samples. While hESC^1q^ equally induce some of the early HEP and CP differentiation markers as hESC^wt^, 68.6% (24/35) and 63.2% (12/19) of HEP and CP lineage-specific markers have a lower expression in the mutant cells (Figure 2H and Figure 2K). HNF4 and GATA4 stainings (Figure 1A, Figure S1) and RNA sequencing data (Figure S4) did not suggest miss-specification in hESC^1q^. While differentiating to HEP and CP, we analysed the expression of markers specific to different stages of the differentiation of these cell types.

Human ESC-derived HEP have a profile between hepatoblast stage 2 and fetal hepatocyte 2 (Figure 2D), and cells with a gain of 1q show overall lower expression of 64% of markers (16/25). Gene-set enrichment analysis of the differential gene expression analysis between HEP^wt^ and HEP^1q^ showed that 145 of the 162 significantly enriched gene-sets in the canonical pathways gene set had negative enrichment scores and that they were frequently related to processes key to hepatocyte and liver function, including cholesterol metabolism, ferroptosis, transsulfuration, plasma lipoprotein remodelling, folate metabolism, selenium micronutrient network and metabolism of steroids (FDR<0.05, Table S2). Gene ontology enrichment analysis showed that 1574 of the 1913 gene sets had negative enrichment scores, including amino acid metabolic process, fatty acid metabolic process and steroid metabolic process (Figure 2J, Table S2).

In the case of CP, CP^1q^ and CP^wt^ equally express a profile between cardiac progenitors and cardiomyocytes, where CP^1q^ have lower expression of genes marking the later stages of differentiation (Figure 2L). Gene set enrichment analysis of the CP samples showed that CAR^1q^ have negative enrichment scores for 157 of the 178 significantly enriched canonical pathway gene-sets, including sets related to dilated cardiomyopathy, folding of actin, striated muscle contraction and hypertrophic cardiomyopathy (Table S2). These genes are key to correct heart contraction functions. Gene ontology enrichment analysis identified negative enrichment scores for 1146 of the 1786 significantly enriched set, including terms, such as l band, Z disc, sarcomere, myofibril, contractile fiber, muscle system process and muscle development (Figure 2M, Table S2).

### Human ESC^1q^ have an *MDM4*-driven competitive advantage that is retained during differentiation

Human ESC^1q^ have a well-established competitive advantage over their genetically balanced counterparts in the undifferentiated state^9,11,14^. We next aimed at determining whether the cells retain this selective advantage during differentiation, and which gene is driving their winning phenotype.

We first looked at the 53 commonly deregulated genes in our HEP, CP and NE samples, and found that *MDM4*, a regulator of p53 activity that is located in the common region of gain and which has been previously suggested as a key gene for the gain of 1q^11^, is consistently upregulated in all samples (Figure 3A). Gene set enrichment analysis of the differentially expressed genes in NE, HEP and CP from 1q and wt cells for the Reactome pathways related to p53 signaling indeed shows that the transcriptional regulation by p53, the regulation of p53 activity, the G1/S damage checkpoint and the p53 dependent responses to DNA-damage in G1 and S are all significantly negatively enriched in cells carrying a gain of 1q (Figure 3B, Table S2, Table S3). This led us to the hypothesis that the higher expression of *MDM4* in cells with a gain of 1q results in the inhibition of p53-mediated transcriptional activation upon DNA damage, leading to a decreased induction of apoptosis upon DNA damage, thus providing a competitive advantage to the mutant cells (Figure 3C).

**Figure 3.**
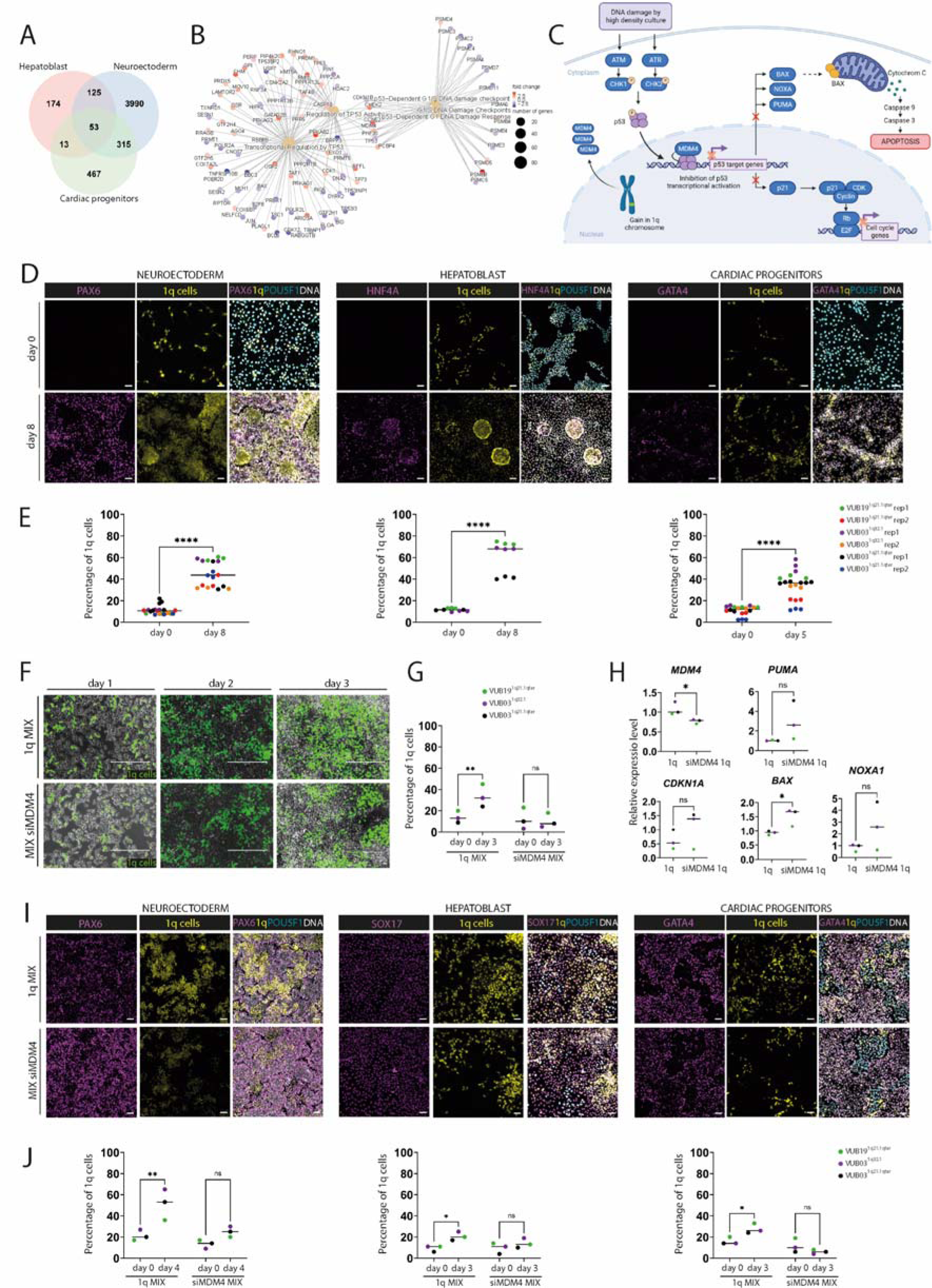
**hESC^1q^ have an *MDM4*-driven competitive advantage that is retained during differentiation** A Venn diagram showing the overlaps in differentially expressed genes between control and 1q-cells, for the three cell types with log_2_fold-change over 0 and FDR<0.05. *MDM4* is part of the core of 53 commonly deregulated genes. B Reactome pathway gene set enrichment analysis of the C2 library canonical pathways related to p53 signaling in NE, HEP and CP from 1q and wt cells. C Graphical summary of the model in which *MDM4* has a higher expression because of the gain of 1q, resulting in an inhibition of the p53 signaling pathway and in turn decreasing the sensitivity of the cells to DNA-damage. D Examples of immunostaining for PAX6, HNF4A and GATA4 after induction cell competition during the differentiation to neuroectoderm, hepatoblast and cardiac progenitors. E Quantification of the numbers of 1q-cells at the start and end of differentiation. F Competition assays in the undifferentiated state. hESC^1q^ are mixed in at a 1:9 ratio with hESC^wt^ and traced by their fluorophore expression. In the si*MDM4* condition, the hESC^1q^ have been treated with siRNA against *MDM4* prior to mixing. G Quantification of the numbers of hESC^1q^ at the start of the cell competition (day 0) and at day 3 of in vitro culture. H mRNA expression of *MDM4* and downstream p53 targets in si*MDM4* treated and untreated hESC^1q^ I Examples of immunostaining for PAX6, SOX17 and GATA4 after induction cell competition during the differentiation to neuroectoderm, hepatoblast and cardiac progenitors, with and without treatment of the hESC^1q^ with siRNA against *MDM4*. J Quantification of the numbers of 1q-cells at the start of the cell competition during differentiation (day 0) and at day 3 or 4, with and without siRNA against MDM4. * p<0.05, ** p<0.01, **** p<0.0001

To determine if cells with a gain of 1q retained their ability to take over the culture during differentiation, we carried out competition assays during NE, HEP and CP induction. For this, 10% of hESC^1q^ stably expressing a fluorescent protein was introduced into a unlabeled hESC^wt^ culture. Differentiation was initiated the next day, and was controlled by immunostaining for PAX6, HNF4A and GATA4 (Figure 3D). To measure culture-take over, the proportion of hESC^1q^ was determined by flow cytometry at the onset and at the end of differentiation.

We found that in all three differentiations, the cells with a gain of 1q outcompeted wild type cells (Figure 3E). During the NE induction, the proportion of 1q cells increased in average 33.9%±2.7% during the 8-day differentiation, from a mean of 10.9% at the onset to 44.9% at day 8 (p <0.0001, unpaired t-test, N=18-21). This was similar for the 8-day differentiation to HEP, where the 1q cells increased in average 49.3%±4.9% (mean at day 0=11.6%, mean at day 8=60.9%, p <0.0001, unpaired t-test, N=9). This increase was less pronounced during the 5-day CP differentiation, with an average 22.7%±3.2% increase (mean at day 0=10.9%, mean at day 5=33.5%, p <0.0001, unpaired t-test, N=18-21), which may be attributable to the shorter time span of the CP differentiation as compared to NE and HEP.

Next, we tested the role of *MDM4* in the competitive advantage of cells with a 1q. For the undifferentiated cells, we downregulated *MDM4* for 24h by siRNA in hESC^1q^ on the day before to start of the competition assay. The cells were imaged daily (Figure 3F), and the proportion of hESC^1q^ was measured with an automated cell counter. On average, hESC^1q^ increased from 14.0 to 33.7% in three days in the untreated condition, while no increase was observed after siRNA treatment (N=3, p^1q^=0.0028, p^si1q^=0.7905, 2-way ANOVA, Figure 3G). For the competition assays during differentiation, the hESC^1q^ were treated for 24h with the siRNA and mixed at a 1:9 ratio, and differentiation was initiated the next day.

Differentiation was shortened to 4 days, as this was the time previous work in the lab showed that a single siRNA transfection could reliably sustain a gene’s downregulation^22^. Figure 3H shows the effect of the siRNA on the expression of *MDM4* and of downstream targets of the p53 signaling pathway in the three hESC^1q^.In the untreated conditions of the competition assays, the fraction of cells with a gain of 1q became significantly larger in all three differentiations (Figure 3I and 3J). The increase was most pronounced after NE induction, with an average increase of 30.0% (p^1q^=0.0053, 2-way ANOVA). In HEP and CP, the mean increases were 11.3% (p^1q^ =0.0131, 2-way ANOVA) and 11.67 (p^1q^ =0.0149, 2-way ANOVA), respectively. Conversely, in the siRNA-treated competition assays, the fraction of 1q cells remained unchanged (Figure 3I and 3J). Taken together, these results show that reducing the levels of *MDM4* in the mutant cells abolishes their competitive advantage both in the undifferentiated state and during differentiation.

### Higher *MDM4* expression in hESC^1q^ results in a decreased sensitivity to DNA-damage induced apoptosis

Next, we sought to elucidate by which mechanisms higher expression of *MDM4* confers the competitive advantage to the cells. We first tested the hypothesis that the higher expression of *MDM4* by cells carrying a gain of 1q leads to a decreased p53-mediated apoptosis in response to DNA damage. For this, we induced DNA damage in hESC^1q^ and hESC^wt^ using bleomycin and carried out a time-course measurement of apoptosis and cell death.

Figure 4A shows the percentages of live and of apoptotic and dead cells for hESC^wt^ (N=3), hESC^1q^ (N=3) and hESC^1q^ treated with siRNA against *MDM4* (N=3), at the start of bleomycin treatment and at the subsequent 2, 4 and 6-hour time-points. hESC^wt^ start undergoing apoptosis at 2h after exposure to bleomycin, followed by a rapid decrease in the numbers of live cells. In contrast, apoptotic cells start appearing in hESC^1q^ as from 4h of exposure and reach 41.9% of apoptotic cells at 6h, as compared to 78.6 % in hESC^wt^ (unpaired t-test, p=0.0169). Treating hESC^1q^ with siMDM4 significantly increases their sensitivity to DNA damage, with apoptosis initiating at 2h, and reaching 49.4% at 4h. siMDM4-treated hESC^1q^ cells do not reach same levels of dead cells at 6h as in hESC^wt^ cells, although the differences are not statistically significant (54.9% vs 78.6%, unpaired t-test, p=0.1530, Figure 4A). This may be explained by an incomplete transfection of the cells, or to overall insufficiently stable downregulation of *MDM4* to the levels of hESC^wt^.

**Figure 4.**
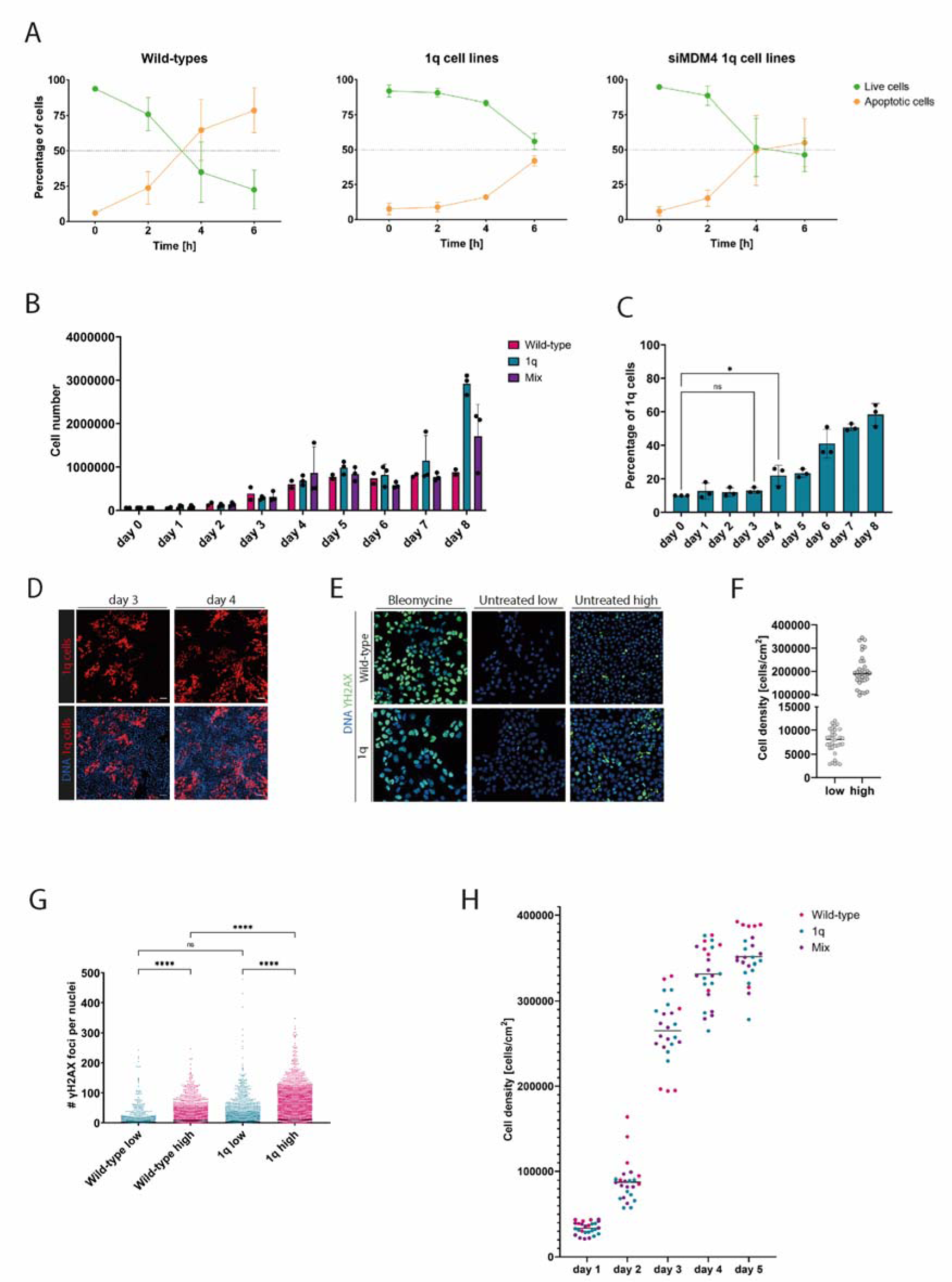
**Higher MDM4 expression in hESC^1q^ results in a decreased sensitivity to DNA-damage induced apoptosis** A Percentages of apoptotic and live cells in hESC^1q^ and hESC^wt^ upon 0h, 2h, 4h and 6h of exposure to bleomycin. B Evolution of the numbers of cells in one culture dish from day 0 to day 8, for hESC^1q^, hESC^wt^ and a 1:9 mix of hESC^1q^ and hESC^wt^. C Percentage of hESC^1q^ over the course of 8 days in culture, starting from a 1:9 mix of hESC^1q^ and hESC^wt^. The proportion of hESC^1q^ becomes significantly increased as from day 4. * p<0.05. D Representative images illustrating the cell density at days 3 and 4 of the time-course experiment. Confluency is reached at day 4 of this experiment. E Examples of the staining for gammaH2AX in hESC^1q^ and hESC^wt^, upon exposure to bleomycin and in low- and high-density cultures. F Cell densities for the low- and high-density culture conditions shown in E and G. The dots represent the counts made in different images of dishes of different cell lines. The line represents the mean of the dots. G Counts of gammaH2AX foci per nucleus in hESC^1q^ and hESC^wt^ at low and high-density culture conditions. Each dot represents a single nucleus, hESC^wt-low^ N=976, hESC^wt-high^ N=2879, hESC^1q-low^ N=2995, hESC^1q-high^ N=7158. **** p<0.0001, one-way ANOVA and Šidák correction.

This raised the question of how a decrease of sensitivity to DNA-damage could provide a competitive advantage to hESC and differentiating cells. We observed during daily monitoring of the competition assays that hESC^1q^ started outcompeting their genetically balanced counterparts once the cultures became confluent. Previous work has indicated that hPSC are prone to replication stress and DNA damage^45,46^, which can be mitigated by addition of nucleosides to the medium^47^ and exacerbated in higher cell density culture due to medium acidification^48,49^. We therefore hypothesized that higher culture cell density generates the conditions for a strong competitive advantage of 1q gains, by increasing the levels of DNA damage.

We studied the cell proliferation dynamics of hESC^wt^ and hESC^1q^ over time in culture in pure form and mixed at a 1:9 ratio (N=3 for each condition). Daily cell numbers count showed that the numbers of hESC^wt^ and hESC^1q^ increased similarly until day 7, suggesting that they have similar cell doubling times. As from day 8, hESC^1q^ continue proliferating even when they have reached a very high density (Figure 4B). Daily analysis of the ratio between hESC^wt^ and hESC^1q^ showed no statistically significant changes until day 3, after which the proportion of hESC^1q^ started steadily increasing (Figure 4C, N=3). This coincided with culture dishes at day 3 still showing empty areas, whereas at day 4 the cells had reached confluence, suggesting that this is a flipping point for 1q gains to start providing a competitive advantage (Figure 4D). To investigate the relationship between cell culture density and DNA damage, we measured DNA damage by γH2AX staining in hESC^1q^ and hESC^wt^, in low and high cell density cultures (mean of 7,547 cells/cm^2^ and 200,050cm/cm^2^ respectively, representative images are shown in Figure 4E and cell counts shown in Figure 4F), and found that cells grown at low density had significantly lower numbers of γH2AX foci than when grown in high density (Figure 4G,wt_low_:11.38, wt_high_:17.95, 1q_low_:13.71, 1q_high_:26.22, p<0.0001, one-way ANOVA). While there were no differences in DNA-damage between in hESC^1q^ and hESC^wt^ grown at low density (Figure 4G, 1.2-fold change, p=0.2314, one-way ANOVA), at high density, hESC^1q^ showed higher γH2AX foci counts than hESC^wt^ (Figure 4G, 1.5-fold change, p<0.0001, one-way ANOVA). Lastly, we counted the numbers of cells in the cell cultures until day 5 and calculated the densities. The mean densities in days 1 and 2 are in the range of the ‘low density’ group in the DNA damage staining, with a mean of 33,711 cells/cm^2^ at day 1 and 87,854 cells/cm^2^ at day 2. From day 3, the densities are in the range and above of the ‘high density’ group, with 265,224cells/cm^2^ at day 3, 331,680 cells/cm^2^ at day 4 and 351,475 cells/cm^2^ at day 5. This indicates that at this point, the cells start undergoing DNA damage more frequently, reaching the condition when a decreased p53-mediated induction of apoptosis starts providing a selective advantage, so cells with a gain of 1q will start outcompeting their genetically balanced counterparts.

## DISCUSSION

Despite that the appearance of abnormal karyotypes in hESC was first noticed nearly 20 years ago^50^ we still have limited insight onto their mechanisms of origin, their driver genes and their functional consequences. In this study, we find that hESC^1q^ have an abnormal response to the differentiation cues to neuroectoderm, with a part of the population miss-specifying to non-CNS cells. Also, while entering the mesendodermal lineage equally efficiently as hESC^wt^, hESC^1q^ generate hepatoblasts and cardiac progenitors that have gene-expression profiles suggestive of a poor cell maturation. It is interesting to note that hESC with a gain of 20q11.21^22,23^, an isochromosome 20^51^ and with a loss of 18q^52^ all display an abnormal response to the neuroectoderm differentiation signaling cues. This common response suggests a particular vulnerability of the neuroectodermal lineage to genetic abnormalities, with potentially common signaling pathways disrupting the differentiation process. In this study, we have identified the alternative cell fate as non-CNS ectoderm, the cells expressing markers of cranial placode and non-neural ectoderm. These cell types are not commonly obtained in the standard NE differentiation by dual-SMAD inhibition, but it has been shown that the addition of different concentrations of BMP4 and modulation the WNT pathway will drive the cells towards these fates^39^. Further, it has been shown that DNA damage-induced stabilization of p53 in hPSC results in an activation of TGF-beta signaling and a downregulation of BMP4 pathway genes^53^, suggesting that deregulation of these pathways specifically in hESC^1q^ during the differentiation process may be at the basis of this alternate fate acquisition.

The differentiation of mesendoderm cell types yielded similar proportions of correctly specified cells, but gene-expression analysis revealed that the hepatoblasts and cardiac progenitors obtained from hESC^1q^ had profiles suggestive of a more immature cell types. Whether this reflects delayed or impaired maturation and whether this extends to other mesendoderm derivates remains to be elucidated, but it further supports the notion that genetic abnormalities do affect the quality of the cells obtained after differentiation, which can negatively impact both the clinical translation of hPSC as well as the reproducibility of results when using these cells in research^54^. Further, here we also show that the genetically variant cells retain their ability to take over a culture during differentiation. Considering that 10 to 20% of cells in an hPSC culture carry a variety of copy number variations^55–57^, and 10% of hESC lines carry mosaic large structural variants^11^, it is possible that the differentiation of a seemingly genetically normal line yields a mixed population of euploid and aneuploid cells with varying degrees of correct cell specification. This lays a further layer of complexity into controlling the outcomes of differentiation, as current methods for genetic screening will easily miss low-grade mosaicism.

In this work, we also identify *MDM4* as the driver gene behind the competitive advantage of the cells with a gain of 1q, and find that this advantage is based on a lower sensitivity to DNA damage-induced apoptosis by the suppression of p53 activity by *MDM4*. An interesting aspect of our findings is that cell density in culture plays a key role in generating the conditions under which hESC^1q^ can outcompete their wild-type counterparts. This finding opens the possibility of tailoring culture conditions to suppress mutant cell take-over, such as by controlling cell density or decreasing DNA damage by regulating the pH of the culture medium^49,56^ or by addition of factors such as nucleosides^58^.

Finally, *MDM4* has recently been shown to also be the driver of the addiction of cancers to chromosome 1q gains to support their malignant growth^59^. The similarities between the genetic abnormalities found in cancers and hPSC have been already noted in the past^60–62^. Gains of 20q11.21, one of the most common acquired abnormalities in hPSC^9^, appear in over 80% of pancreatic and colorectal carcinomas ^63,64^ and gains of 1q appears in over half of the cases of hepatocellular carcinoma^65^ and have been recently identified as the first copy number alterations occurring in breast cancer and melanoma evolution^59^. These parallels, together with the high incidence of the gain of 1q in culture, the fact that the variant cells can take over also during differentiation and that the gain of 1q impacts the final cell product, stress the importance of factoring in the genetic integrity of the cells not only in a clinical setting, but also in a research context to ensure reliable and reproducible science^66,68^.

## MATERIALS AND METHODS

### hESCs lines, cell culture and banking and transgenic modification

All the hESC lines were derived from human embryos and characterized as described in ^26,27^. They are registered in the EU hPSC registry (https://hpscreg.eu/). VUB03 and VUB19 remained genetically balanced up to passage 20 and 24, respectively. All stocks of hESC lines are preserved in liquid nitrogen in 90 % Knock-out serum (Thermo Fisher Scientific) and 10% sterile DMSO. hESC were cultured in dishes coated with human recombinant Laminin-521 (Biolamina). Laminin-521 was diluted in phosphate buffered saline (PBS) (Thermo Fisher) with calcium and magnesium. Coated dishes were stored overnight at 4°C before the use. NutriStem™ (Biological Industries) medium with 10mM penicillin/streptomycin was changed daily. hESC cultures were kept in the incubator at 37°C and 5% CO2. For passaging, cells were washed with PBS and incubated with recombinant enzymes TrypLE (Thermo Fisher) for 10 minutes at 37°C for single cell dissociation. TrypLE was deactivated with NutriStem™ medium and cells were centrifuged at 1000 rpm for 5 min. Pellet was resuspended in 1 ml of medium and counted with image-based cytometer Tali™ (Invitrogen) if needed or transferred in a new dish in ratio between 1:20 to 1:50.

Labelled hESC1q cells were generated by infecting hESC1q with lentiviral particles expressing fluorescent protein. HEK293T cells were transfected with pMDG (VSV.G encoding plasmid), pCMVΔR8.9 (gag---pol encoding plasmid) as packing vectors and three different vectors each with their own fluorescent protein (LeGO---EF1a---V2---Puro (Venus fluorescent protein), LeGO---EF1a---C2---Puro (mCherry) and pCDH---EF1---MCS---pA---PGK---copGFP---T2A---Puro (Green fluorescent protein). Vectors with mCherry and Venus fluorescent protein were kindly provided by Kristoffer Riecken^67^. Vector with GFP was a gift from VUB Laboratory for molecular and cellular therapy. Vector was purchased from Addgene (https://www.addgene.org/17448/) and cloned CMV promoter to EF1a. Lentiviruses were produced by transfection of HEK 293T cells with VSV.G and gag-pol plasmids together with the plasmid of interest and PEI (2ug per ug of DNA, Polysciences Inc) in Opti-MEM medium (Thermo Fisher Scientific). After 4h of transfection the transfection cocktail was replaced by complete medium. Supernatant containing lentiviral particles was collected after 48 and 72h and stored at -80C. A day before hPSC were seeded at the density 50.00cells/cm2. hESC were transduced with 1:1 mix of Nutristem and complete medium containing lentivirus together with 1:1000 protamine sulfate (LEO Pharma; 10mg/ml). Cells were incubated with transduction cocktail for 4h. Next, cells were washed 5 times with PBS and refreshed with Nutristem medium. Selection of successfully transduced cells was done with FACS.

Working cell banks comprising labeled 1q cell lines and their isogenic counterparts were established, which were additionally controlled for cell identity and karyotype. We utilized the working cell bank for subsequent experiments and the cells were not kept in culture for more than 5 passages after thawing to prevent them from genetic drift.

### Genome characterization

Genomic DNA was extracted with DNeasy Blood & Tissue Kit (Qiagen) following producer’s instructions. Concentration was assessed with NanoDrop 1000 (Thermo Fisher Scientific). DNA was stored at +4°C. The genetic content of the hESCs was assessed through shallow whole-genome sequencing by the BRIGHTcore of UZ Brussels, Belgium, as previously described^69^. Copy-number assays were used to control for gains of 1q, 12p and 20q11.21 at regular intervals. Real time polymerase chain reaction was performed on ViiA™ 7 system (Applied Biosystems). Total volume of qPCR reaction was 20µl and it contained 10µl of 2x qPCR Master Mix Low ROX (Eurogentec), 1µl of 20x TaqMan Copy Number Assay (Life Technologies), 1µl of nuclease-free water and 40 ng of DNA. TaqMan™ Copy Number Reference Assay RNase P (Applied Biosystems, 4403328) was added instead of nuclease-free water. Genomic DNA from leukocytes of healthy donor was used as a control. Non template control was tested in each experiment. Cycling parameters are 2min at 50°C, 40 cycles with 10min at 95°C and 15 sec at 95°C and 1 min at 60°C. Each sample was tested in triplicates. Copy number assays used were KIF14 (Thermo Fisher, Hs_cn), NANOG (Thermo Fisher, Hs_cn) and ID1(Thermo Fisher, Hs_cn). Experiments were run as either comparative Ct (cycle threshold) or relative standard curve analysis. Data analysis was performed by ViiATM 7 v2.0 software or Copy Caller v2.1 (Applied Biosystems).

### Neuroectoderm differentiation

The protocol for neuroectoderm induction was adapted from Douvaras et al., 2015^70^. We seeded 22.500 cells/cm2 on dishes coated with Laminin-521 and 1:100 RevitaCell (Thermo Fisher Scientific). After 24h differentiation was induced with freshly preparred neural induction medium containing from DMEM/F12 (Thermo Fisher Scientific), 1x NEAA (TFS), 1x GlutaMAX (TFS), 1x 2-mercaptoethanol (TFS), 25 ug/ml insulin (Sigma-Aldricht) and 1x penicillin/streptomycin (TFS). The medium was supplemented with 10μM SB431542 (Tocris), 250 nM LDN193189 (STEMCELL Technologies) and 100 nM RA (Sigma-Aldrich). The medium was refreshed every day for 8 days.

### Cardiac progenitor differentiation

The protocol for cardiac progenitor differentiation was modified from Lin et al., 2020^71^. hESC were seeded at the density of 22.500 cells/cm2 on Laminin-521 coated plates. RevitaCell was added to the cell suspension in 1:100 dilution before the seeding. When cells reached 90% confluency, differentiation was initiated with cardiomyocyte differentiation basal medium (CDBM) supplemented with 5µM CHIR99021 (Tocris). After 24h treatment with CHIR99021, medium was changed with fresh CDBM with addition of 0,6U/ml heparin (Sigma-Aldricht). For the next 3 days CDBM with 0,6U/ml heparin and 3mM IWP2 (Tocris) was refreshed daily. CDBM medium was composed of DMEM/F12 (Thermo Fisher Scientific)), 64 mg/L L-ascorbic acid (Sigma-Aldrich), 13.6 ug/L sodium selenium (Sigma-Aldrich), 10ug/ml transferrin (Sigma-Aldrich) and 1x chemically defined lipid concentrate (Thermo Fisher Scientific)).

### Hepatoblast differentiation

The hepatoblast differentiation protocol was based on protocol from Boon et al. 2020^72^. hESC were seeded with 1:100 RevitaCell (Thermo Fisher Scientific)) at the density of 35.000cells/cm2 on Laminin-521 coated plates. Next day Nutristem medium was refreshed. Differentiation was initiated the next day with liver differentiation medium (LDM) supplemented with 50 ng/ml Activin A (STEMCELL Technologies), 50ng/WNT3A (PreproTech) and 6ul/ml DMSO (Sigma-Aldrich). After 48h, medium was refreshed without WNT3A factor. On days 4 and 6 LDM medium with added 50ng/ml BMP4 (STEMCELL Technologies) and 6ul/ml DMSO was used. LDM medium was composed of MCDB 201 medium with pH 7,2 (Sigma-Aldrich), DMEM High Glucose medium (Westburg Life Sciences), L-Ascorbic Acid (Sigma-Aldrich), Insuli-Transferrin-Selenium (ITS-G, Thermo Fisher Scientific), Linoleic Acid-Albumin (LA-BSA, Sigma-Aldrich), 2-Mercaptoethanol (Thermo Fisher Scientific) and Dexamethasone (Sigma-Aldrich).

### Total RNA isolation, cDNA synthesis and quantitative real time PCR

RNA was extracted from cell pellets with RNeasy Mini Kit (Qiagen) following the producer’s protocol. Concentration of obtained RNA was measured with NanoDrop 1000 (Thermo Fisher Scientific). RNA was stored at -80°C. Reverse transcription of RNA to cDNA was performed with First-Strand cDNA Synthesis Kit (GE Healthcare) following producer’s instructions. cDNA was stored at -20°C.

Real-time qPCR was performed on ViiA™ 7 system (Applied Biosystems). Total volume of qPCR reaction was 20µl and it contained 10µl of 2x qPCR Master Mix Low ROX (Eurogentec), 1µl of 20x TaqMan Gene Expression Assay (Life Technologies), 1µl of nuclease-free water and 40 ng of cDNA. Each sample was tested in triplicates, GUSB was used as a house-keeping gene. Genes and assays are listed in Table S4. We used the standard cycling parameters of the Viia7 instrument. Experiments were run as comparative Ct (cycle threshold) analysis. Data analysis was performed by ViiATM 7 v2.0 software.

### Immunofluorescent Staining

Cells were washed 3 times with PBS and incubated for 15 min with 3,6% paraformaldehyde at room temperature for fixation. Cells were washed again 3 times with PBS. Blocking was performed with 1-2 hour incubation with 10% fetal bovine serum (FBS). Primary antibodies were diluted in specific ratio in 10 % FBS and incubated with cells for 1 hour in the dark at room temperature. After washing 3 times with PBS, secondary antibodies were diluted in FBS together with 1:2000 dilution of Hoechst 33342 (Life Technologies) and incubated for 2 hours at the room temperature. Cells were again washed three times with PBS and stored at 4°C until analyzed with confocal microscope Zeiss. List of antibodies is shown in Table S4.

### Flow cytometry and Annexin V staining

The hESC were harvested by incubating them 10 min at 37°C with 1 ml of TrypLE (Thermo Fisher). The cell suspension was gently pipetted up and down with 2 ml of medium and strained through 20 µm cell strainer (pluriStrainer) to eliminate any cell clumps. Single cell suspension was centrifuged at 1000 rpm for 5 min. Medium was aspirated, cells were resuspended in 1 ml of PBS and spun down at 1000 rpm for 5 min. If needed, the cells were incubated with Alexa Fluor® 647 Annexin V (BioLegend) based on the manufacturer’s instructions. Next, the cells were washed with 1mL of PBS and centrifuged. Incubation with Live/dead stain (Thermo Fisher Scientific) was performed following the producer’s protocol. Cells were washed again and fixed with 3,6% paraformaldehyde at room temperature for 10 min. After centrifugation (1000 rpm for 5 min) the paraformaldehyde was aspirated, the cells were washed with 1 ml of PBS and spun down. The fixed cells were resuspended in PBS and stored at 4°C until the analysis with Flow cytometer Aria III (BD).

### RNA Sequencing

RNA-seq library preparation was performed using QuantSeq 31 mRNA-Seq Library Prep Kits (Lexogen) following Illumina protocols. Sequencing was performed on a high-throughput Illumina NextSeq 500 flow cellThe FastQC algorithm ^73^ was used to perform quality control on the raw sequence reads prior to the downstream analysis. The raw reads were aligned to the new version of the human Ensembl reference genome (GRCh38.p13) with Ensembl (GRCh38.83gtf) annotation using STAR version 2.5.3 in 2-pass mode^74^.The aligned reads were then quantified, and transcript abundances were estimated using RNA-seq by expectation maximization (RSEM, version 1.3.3)^75^.

The count matrices were imported into R software (version 3.3.2) for further processing. The edgeR^76^ package was utilized to identify differentially expressed genes (DEGs) between groups. Transcripts with a count per million (cpm) greater than 1 in at least two samples were considered for the downstream analysis. Genes with a log2-fold change greater than 1 or less than -1 and a false discovery rate (FDR)-adjusted P-value less than 0.05 were considered significantly differentially expressed. Volcano plots of DEGs were generated using the ggplot2^77^ package in R.

Principal component analysis (PCA) and heatmap clustering were performed using normalized counts and R packages. The heatmap was generated using the heatmap.2 funtion. PCA was performed using the prcomp function and plotted using ggplot2. Gene set enrichment analysis (GSEA), gene ontology enrichment analysis and reactome pathway gene set enrichment analysis was applied to detect the enrichment of pathways using the fgseaMultilevel, gseGO and enrichPathway function in R. Values are ranked by sign(logFC)*(-log10(FDR)). |NES| > 0 and adjusted p-value < 0.05 were considered the thresholds for significance for the gene sets.

### Downregulation of MDM4 gene with siRNA

SMARTpool siRNA for MDM4 was purchased from Horizon Discovery. 1q hESC were seeded at the density 40.000 cells/cm2. After 24h cells were transfected with 50nM SMARTpool siRNA in Nutristem medium without antibiotics together with RNAiMAX (Thermo Fisher Scientific), according to manufacturer’s instructions. Transfection cocktail was added to the cells 24h prior to seeding for differentiation.

### Statistics

All differentiation experiments were carried out in at least triplicate (n ≥ 3). All data are presented as the mean ± standard error of the mean (SEM) or standard deviation (SD). Statistical evaluation of differences between 2 groups was performed using unpaired two-tailed t tests, one-way or two-way ANOVA in GraphPad Prism9 software, with p < 0.05 determined to indicate significance.

### Data availability

The RNA sequencing counts per million tables are provided in the supplementary material. The raw sequencing data are available upon request. Source data can be retrieved from osf.io/qzyp4.

## Author contributions

N.K. carried out all of the experiments and bioinformatics analysis unless stated otherwise and co-wrote the manuscript. E.C.D.D. assisted with the bioinformatics analysis. Y.L., and D.AD. assisted with the cell culture. S.V and L.A.vG assisted with the hepatoblast differentiation and the flow cytometry K.S. edited the paper. C.S. cowrote the manuscript and designed and supervised the experimental work. All co-authors provided scientific input and proof-read the manuscript.

## Acknowledgments

The authors wish to thank Kristoffer Riecken of the University of Hamburg for kindly sharing the lentiviral constructs for the overexpression of the fluorescent proteins mCherry and Venus. Thanks also go to Yannick De Vlaeminck for the assistance with production of lentivirus and kindly sharing the modified GFP plasmid. Y.L. is a predoctoral fellow supported by the China Scholarship Council (CSC), and N.K., M.R. and E.C.D.D. are predoctoral fellows supported by the Fonds voor Wetenschappelijk Onderzoek Vlaanderen (FWO). This research was supported by the Methusalem Grant to Karen Sermon (Vrije Universitet Brussel).

