## supplemental data for "Gain of 1q confers an MDM4-driven growth advantage to undifferentiated and differentiating hESC while altering their differentiation capacity"

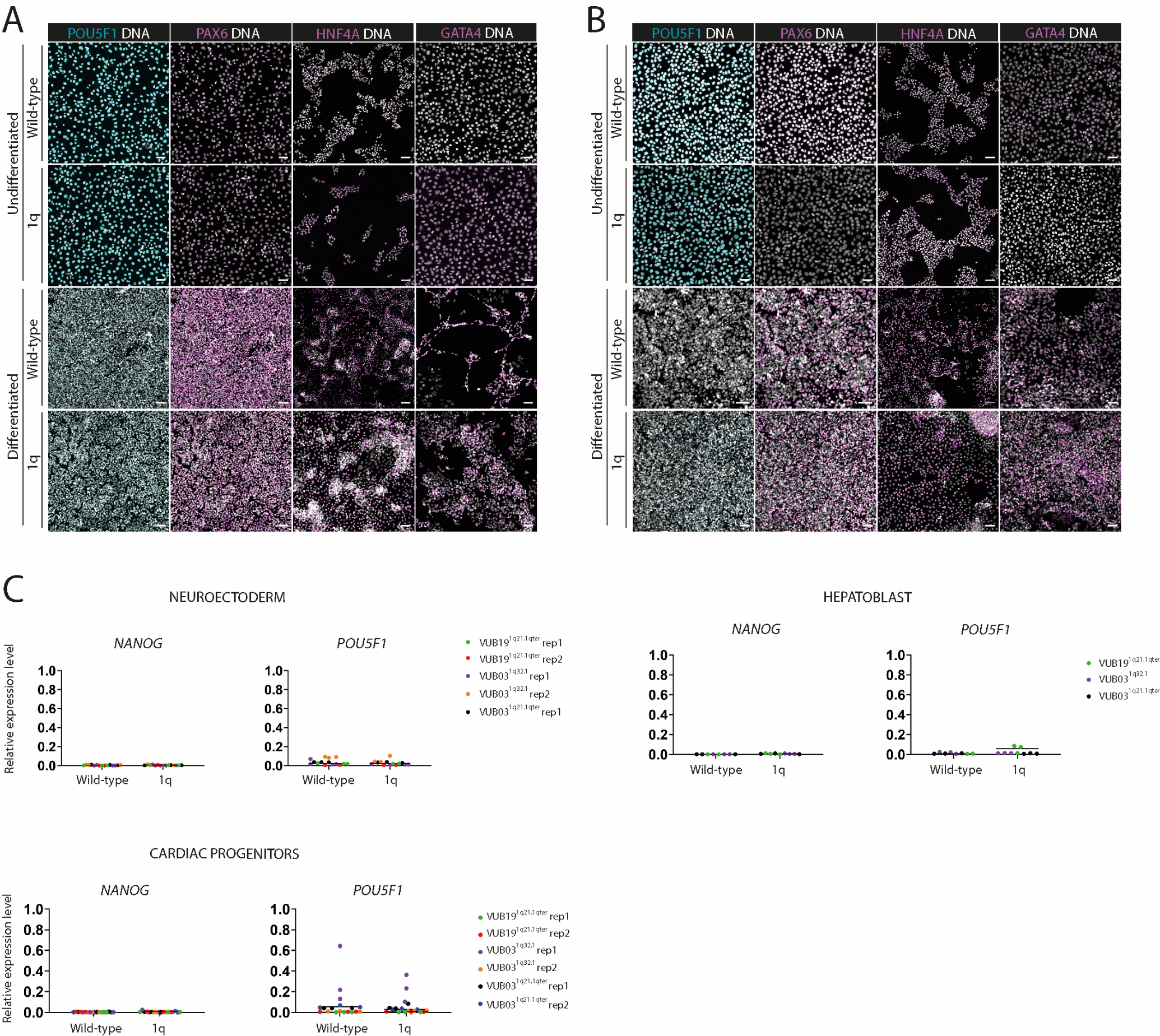

**Figure S1. Differentiation of hESC to neuroectoderm, hepatoblast and cardiac progenitors results in downregulation of pluripotency markers and upregulation of differentiation markers in wild-type and 1q cells.**

**A** Immunostaining of pluripotency marker POU5F1 and neuroectoderm marker PAX6, hepatoblast marker HNF4A and cardio progenitor marker GATA4 of cell lines VUB03^wt^, VUB03^1q32.1^. Markers are shown in wild-type cell line and their 1q counterparts before and after the differentiatiation.

**B** Immunostaining in cell lines VUB03^wt^ and VUB03^1q21.1qter^.

**C** Expression of pluripotency markers NANOG and POU5F1 determined with qPCR in two wild-type and three 1q cell lines. Expression is shown relative to wild-type hESC. Each independent experiment replicate (rep) is coded with the unique color.

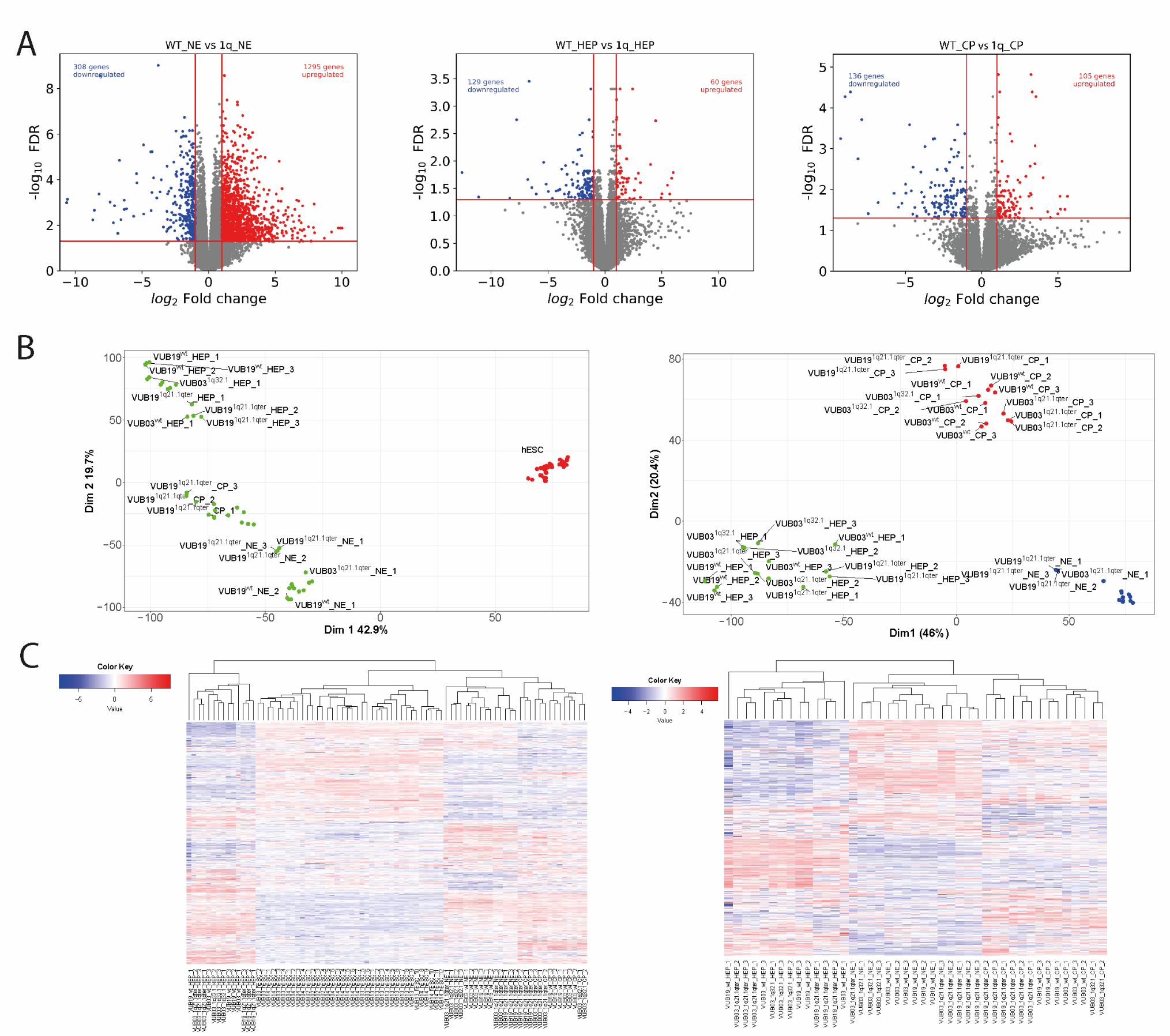

**Figure S2. Volcano plots, PCA and unsupervised heatmaps of the mRNA sequencing of NE, CP and HEP from hESC^wt^ and hESC^1q^**

**A** Volcano plots represent differentially expressed genes after neuroectoderm, hepatoblast and cardiac progenitor differentiation. Results are shown as 1q cells relative to wild-type differentiated cells. Significant results are considered at |log_2_ fold change|>1 and FDR< 0.05. In neuroectoderm differentiation 308 genes are downregulated and 1295 upregulated, in hepatoblast 129 downregulated and 60 upregulated genes and in cardiac progenitor cells 136 downregulated and 105 upregulated genes.

**B** PCA results of wild-type and 1q cell lines differentiated to 3 germ layers in relation to hPSC (left) and differentiated wild-type cells in relation to differentiated 1 q cell lines. Cell lines from the same lineage are clustering together.

**C** Unsupervised heatmap of all differentially expressed genes in differentiated and hESC lines (left) and only differentiated cell lines (right).

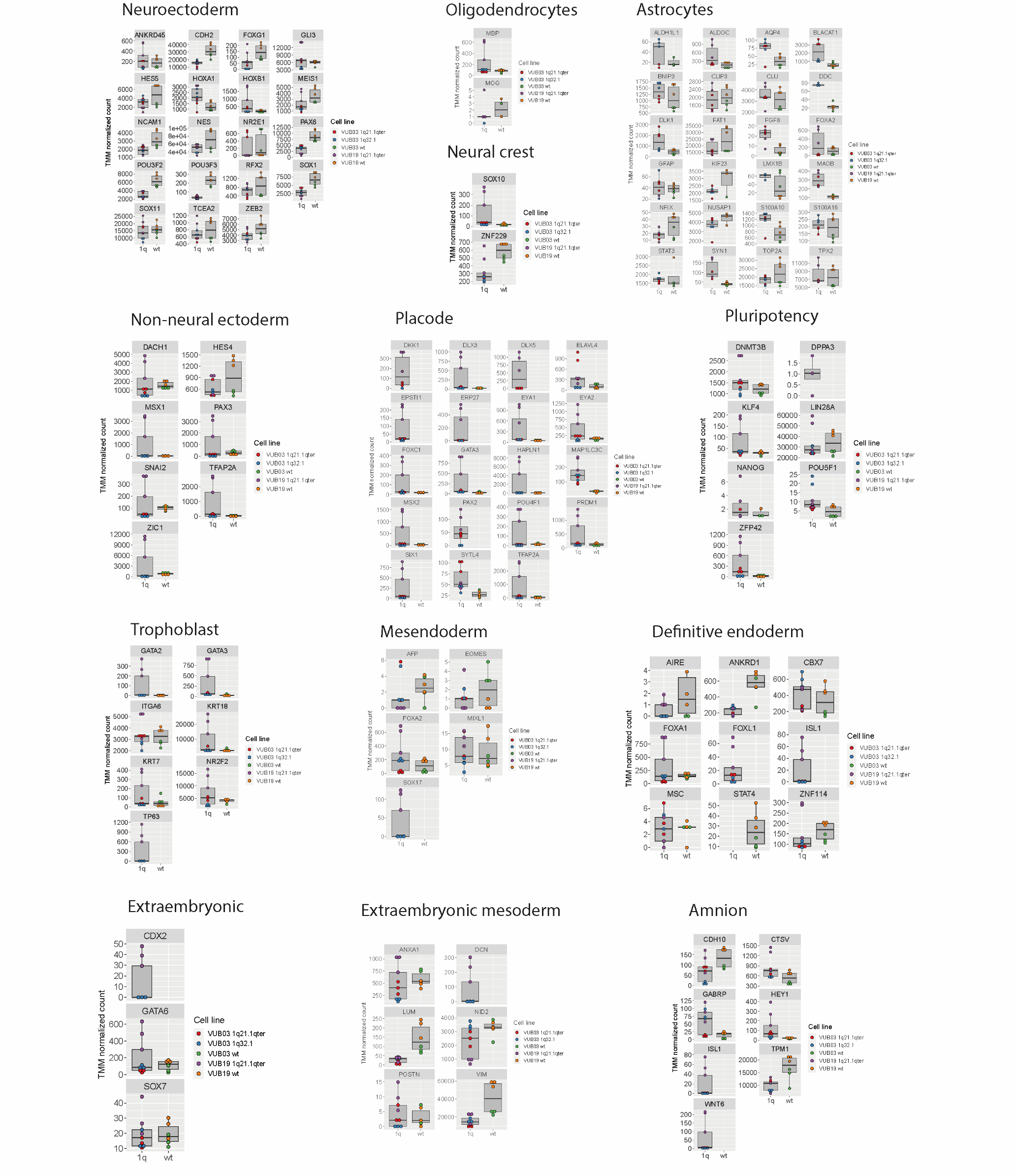

**Figure S3. Box plot of TMM normalized counts per gene in individual cell lines of neuroectoderm differentiated wild-types and 1q cell lines. Expression of different cell type markers is represented.**

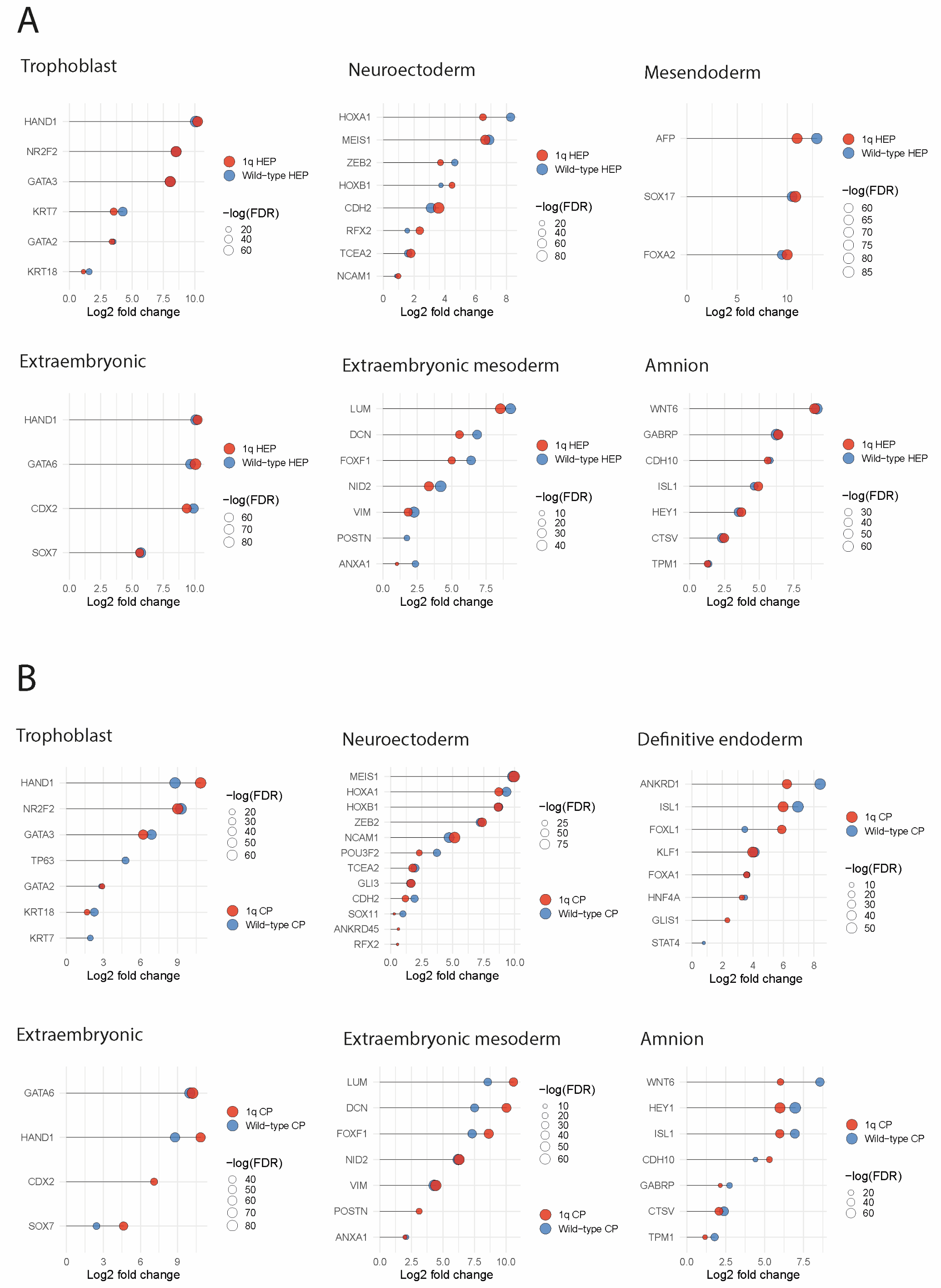

**Figure S4. Studying miss-specification in hepatoblast (A) and cardiac progenitor (B) wild-type and 1q cell lines. Lollipop plots showing log_2_ fold change expression of different cell type markers.**
